# Healthy Ageing and Sentence Production: Disrupted Lexical Access in the Context of Intact Syntactic Planning

**DOI:** 10.1101/327304

**Authors:** Sophie M. Hardy, Katrien Segaert, Linda Wheeldon

## Abstract

Healthy ageing does not affect all features of language processing equally. In this study, we investigated the effects of ageing on different processes involved in fluent sentence production, a complex task that requires the successful execution and coordination of multiple processes. In Experiment 1, we investigated age-related effects on the speed of syntax selection using a syntactic priming paradigm. Both young and older adults produced target sentences quicker following syntactically related primes compared to unrelated primes, indicating that syntactic facilitation effects are preserved with age. In Experiment 2, we investigated age-related effects in syntactic planning and lexical retrieval using a planning scope paradigm: participants described moving picture displays designed to elicit sentences with either initial coordinate or simple noun phrases and, on half of the trials, the second picture was previewed. Without preview, both age groups were slower to initiate sentences with larger coordinate phrases, suggesting a similar phrasal planning scope. However, age-related differences did emerge relating to the preview manipulation: while young adults displayed speed benefits of preview in both phrase conditions, older adults only displayed speed preview benefits within the initial phrase (coordinate condition). Moreover, preview outside the initial phrase (simple condition) caused older adults to become significantly more error-prone. Thus, while syntactic planning scope appears unaffected by ageing, older adults do appear to encounter problems with managing the activation and integration of lexical items into syntactic structures. Taken together, our findings indicate that healthy ageing disrupts the lexical, but not the syntactic, processes involved in sentence production.

## General Introduction

Producing a fluent and coherent sentence is a complex task involving the coordination of multiple cognitive and neural mechanisms (Levelt, 1989; Mody, 2017). As we age, changes occur that can create challenges for language processing, such as a widespread reduction in grey matter volume (Good et al., 2001) and a decline in working memory capacity (Waters & Caplan, 2003). Nevertheless, older adults have a wealth of experience with language and are often able to adopt effective processing strategies, such as the recruitment of addition brain areas, to compensate for lost efficiency elsewhere (Reuter-Lorenz & Park, 2014; Wingfield & Grossman, 2006). This paints a multifactorial picture of language processing in old age in which some language skills decline because of age-related cognitive changes, but in which others are preserved because of the successful adoption of compensation strategies (for reviews, Burke & Shafto, 2008; Peelle, 2019). Investigating how different aspects of language processing are affected by old age is critical for understanding this complex balance between decline and preservation. In this study, we conducted two novel experiments investigating age-related changes in sentence production; specifically, we investigated the processes involved in syntax generation (Experiment 1), as well as sentence planning and lexical retrieval (Experiment 2). Our findings reveal a contrast between the preservation of syntactic skills, but the disruption of lexical access, in old age; this adds to the growing evidence that healthy ageing does not affect all features of language processing to the same extent.

A number of previous studies have demonstrated age-related decline in language production. To first consider age-related changes at the word level, several studies have found older adults to be slower and more error-prone in picture naming tasks, particularly for low-frequency words (see Feyereisen, 1997, for a review), and to experience increased tip-of-the-tongue states in which a speaker is certain that they know a word but is unable to produce it (Burke, MacKay, Worthley, & Wade, 1991; Segaert et al., 2018; Shafto, Burke, Stamatakis, Tam, & Tyler, 2007). This suggests an increased difficulty in retrieving the name of a lexical object and its corresponding phonological form, something which may be attributable to age-related atrophy in the left insula (Shafto et al., 2007). Age-related deficits are also found at the sentence level of production: with age, there is a decline in the production of complex syntactic structures, such as embedded clauses, coupled with an increase in syntactic errors, such as the use of the incorrect tense (Kemper, 1987; Kemper, Greiner, Marquis, Prenovost, & Mitzner, 2001; Kemper, Herman, & Liu, 2004; Kemper & Sumner, 2001; Rabaglia & Salthouse, 2011). This apparent decline in syntax production is often considered to arise from age-related decreases in the capacity or efficiency of working memory, a cognitive resource that is critical when producing complex sentences that contain multiple clauses and that require greater syntactic operations of movement (Abrams & Farrell, 2011; Kemper & Sumner, 2001; MacDonald & Christiansen, 2002).

In contrast, other aspects of language production are characterized by stability and even improvement with age. Most notably, vocabulary size and knowledge consistently increase with age (Verhaeghen, 2003). Older adults also appear to perform similarly to young adults in tasks where they must switch between formulating alternative syntactic structures, such as dative verb and transitive verb alternatives (Altmann & Kemper, 2006; Davidson, Zacks, & Ferreira, 2003). Moreover, in situations in which the task demands are reduced, minimal age differences are found; for example, Kemper, Herman, and Lian (2003) found that young and older adults produced similar responses when asked to incorporate intransitive (‘smiled’) or transitive (‘replaced’) verbs into their sentences, and age differences in fluency only emerged when participants were asked to incorporate more complex complement-taking verbs (‘expected’). This effect of task complexity on language production skills in old age can be best explained by Peelle’s (2019) ‘supply and demand’ framework, which suggests that behavioural success reflects a complex balance between specific task requirements and the level of cognitive resources available to the speaker; specifically, if task requirements outweigh cognitive resources, processing efficiency will decline, leading to poor performance. Due to overall neuroanatomical and cognitive changes that occur during healthy ageing, it is no surprise that older adults’ neurocognitive capacity for any given language task is likely to be less than young adults. However, this does not necessarily mean that age differences will always emerge: older adults may still perform similarly to young adults when task requirements are sufficiently low (e.g., when producing simpler syntactic constructions) or they may adopt compensatory processing strategies (e.g., the recruitment of other brain areas). In this way, identical behavioural performance in young and older adults may not always reflect identical neural or cognitive processes.

The idea of neural compensation in ageing has been studied most in terms of language comprehension in brain imaging studies that have demonstrated that older adults engage additional brain areas in order to maintain high levels of accuracy (see Wingfield & Grossman, 2006, for a review). Likewise, older adults may employ different strategic approaches in order to compensate for processing deficits elsewhere, such as a greater reliance on discourse during reading (Stine-Morrow, Miller, Gagne, & Hertzog, 2008). These same principles of compensation can also be applied to production; for example, Altmann and Kemper (2006) suggested that the minimal age group differences they observed in their sentence generation task were the result of older adults adopting a different strategy to young adults (they always assigned the top-presented lexical item to the subject role when producing a sentence to describe a multi-word display). Overall, this highlights the importance of continuing to investigate the effect of ageing on different aspects of language processing. Moreover, even when there appear to be no group differences, this does not necessarily mean that young and older adults are engaging the exact same processing networks.

The aim of our study was to investigate how the syntactic and lexical processes involved in sentence generation are affected by healthy ageing using paradigms that have not previously been used with older adults. In both experiments, we employed on-line onset latency measures of sentence production in order to gain information about the incremental fashion in which sentences are planned and produced (see, Wheeldon, 2013, for a review of latency measures of speech production). Most previous studies investigating sentence production and ageing have predominantly used off-line measures, involving the assessment and coding of sentences after they have been produced (e.g., Kemper et al., 2001, 2003, 2004; Rabaglia & Salthouse, 2011), which, while informative about syntactic choices and errors, cannot provide insight into the time-course of the underlying sentence generation process (Marinis, 2010; Mertins, 2016). To our knowledge, only a handful of studies to date have investigated older adults’ sentence production using on-line measures (Griffin & Spieler, 2006; Spieler & Griffin, 2006); hence, there remains a considerable gap in the ageing literature regarding the timing of speech preparation and how different syntactic and lexical processes unfold during the course of sentence production.^1^ In Experiment 1, we used a syntactic priming paradigm (as in Smith & Wheeldon, 2001; Wheeldon & Smith, 2003) to investigate age-related differences in the speed of syntax generation. In Experiment 2, we used a planning scope paradigm with an embedded picture preview element (as in Smith & Wheeldon, 1999; Wheeldon, Ohlson, Ashby, & Gator, 2013) to investigate age-related differences in syntactic planning scope and lexical retrieval. Taken together, the two experiments provide insight into the effect of healthy ageing on critical features of sentence production that must all be executed quickly and efficiently for a speaker to produce a fluent and coherent sentence; specifically, we focus on syntax generation (Experiment 1) and initial sentence planning (Experiment 2).

## Experiment 1

### Examining the Effect of Ageing on Latency Measures of Syntax Facilitation

The process of producing a sentence begins with the preparation of a preverbal message – this is a conceptual representation of all the information that the speaker wishes to convey and that will ultimately be formulated into a coherent grammatical structure (Levelt, 1989; Levelt, Roelofs, & Meyer, 1999). The exact structure of preverbal messages is debated, but it is generally agreed that they minimally contain conceptual category information and a thematic structure with concepts assigned to thematic roles (Wheeldon, 2013). The preverbal message triggers the formulation stage in which the message is turned into linguistic representations, involving both the rapid retrieval of lexical items and the generation of an appropriate syntactic structure, which must be integrated correctly to convey the intended message. More traditional models of sentence production propose that grammatical encoding is lexically driven such that lemmas (representations of the syntactic and semantic properties of a word) are first selected and assigned grammatical roles (e.g., subject or object), which then drive the generation of a syntactic structure (Bock & Levelt, 1994; Levelt et al., 1999; Pickering & Branigan, 1998). Alternatively, computational models postulate that there is a complete dissociation between syntax generation and lexical retrieval such that syntactic structure is derived solely from conceptual structure (i.e., thematic roles) with lexical access occurring independently (Chang, Dell, & Bock, 2006; Chang, Dell, Bock, & Griffin, 2000).

While there remains debate about the exact relationship between syntax generation and lexical retrieval (see Wheeldon, 2011, for a review of the evidence for both lexically mediated and lexically independent models), it is widely agreed that sentence production occurs incrementally such that only a small amount of planning occurs prior to articulation and that planning continues to unfold after speech onset for the remainder of the sentence (Levelt, 1989, 1992). Consequently, the amount of time that a speaker takes to begin a sentence is informative about the amount of planning that has occurred prior to speech onset in terms of both the retrieval of lexical items and the generation of syntax (Levelt, 1989; Wheeldon, 2013). On-line onset latency measures can therefore be used to explore age-related differences in the type and amount of advanced planning, or scope, of the sentence generation process.

One paradigm that has been used to explore the processes involved in syntax generation is *syntactic priming*. Broadly speaking, syntactic priming refers to the facilitation of syntactic processing that occurs when a syntactic structure is repeated across an otherwise unrelated prime and target (Bock, 1986; Pickering & Ferreira, 2008). *Choice syntactic priming* is the phenomenon whereby speakers are more likely to repeat a syntactic structure that they have recently processed (see Mahowald, James, Futrell, & Gibson, 2016, for a meta-analytical review). In our study investigating the speed of syntax generation, we were interested in *onset latency syntactic priming:* the facilitated speed of syntactic processing that occurs when a syntactic structure is repeated across a prime and a target (Corley & Scheepers, 2002; Segaert, Menenti, Weber, & Hagoort, 2011; Segaert, Weber, Cladder-Micus, & Hagoort, 2014; Segaert, Wheeldon, & Hagoort, 2016; Wheeldon & Smith, 2003).

For example, using a picture description task, Smith and Wheeldon (2001) demonstrated that when a speaker must produce a given syntactic structure on a target trial (1a), this was initiated quicker (i.e., decreased speech onset latencies) following recent production of the same structure (1b), compared to when a different structure had just been produced (1c).

**(1a)** Target: “the spoon and the car move up”

**(1b)** Related prime: “the eye and the fish move apart”

**(1c)** Unrelated prime: “the eye moves up and the fish moves down”

This latency priming effect cannot have its source in conceptualization, lexical access or phonological planning as these factors were tightly controlled within the experimental design (i.e., there was no prosodic, visual or lexical similarity between any of the corresponding primes and targets). Further experiments by Smith and Wheeldon (2001) also ruled out alternative explanations relating to overall sentence complexity (the effect persists when both the related and unrelated prime feature the same number of clauses as the target), as well as to visual perception and picture movement (the effect persists when the related and unrelated primes feature the exact same movement patterns, and when stationary written prime sentences are used). This indicates that the facilitation effect observed is specifically related to the repetition of syntactic structure between the prime and target. Similar facilitation effects have been observed during sentence comprehension, as evidenced by reduced reading times when a structure is repeated between the prime and target (Tooley, Traxler, & Swaab, 2009).

The two most common theoretical accounts of structural priming relate to the residual activation of a prime syntactic structure (Pickering & Branigan, 1998) and implicit learning processes that occur when an unexpected prime is heard (Chang et al., 2006). However, these models only provide explanations of facilitation effects relating to syntactic choices and not to the speed of sentence production; thus, the models offer minimal insight into the mechanisms that underlie onset latency syntactic priming. By contrast, Segaert et al. (2016) proposed a two-stage competition model that explains the effect of syntactic priming on both choices and onset latencies (see also Segaert et al., 2011, 2014). According to the model, alternative syntactic structures (e.g., active vs. passive) are represented by syntactic nodes that transmit activation and inhibition (i.e., negative activation) to neighbouring nodes within the network (i.e., to the competing syntactic alternative). The activation levels of each node, and thus how much inhibitory activation is transmitted to the competing node, are determined by the relative frequency of the structure (established through implicit learning). Sentence production begins with construction of the preverbal message and this is followed by two sequential stages. First is the selection stage during which a speaker selects one syntactic structure from competing alternatives. Next follows the planning stage during which the selected syntax is incrementally planned and produced. While syntactic choice is determined solely at the selection stage, production speed is determined by the additive time taken to complete both stages.

Consequently, in a task in which the choice element is removed and there are very clear instructions about which syntactic structures to produce (as in Smith & Wheeldon, 2001), speech onset latencies are largely determined by processing at the planning stage with very minimal processing required at the selection stage as there are no competing syntactic alternatives. This is because a high level of certainty about what sentence types to use and when to use them means that there is very little (if any) syntactic choice involved prior to sentence planning. This is in contrast to a choice syntactic priming task, in which participants are typically asked to describe pictures that could be described using multiple syntactic alternatives (Pickering & Ferreira, 2008). In this study, we therefore investigated age-related effects on onset latency syntactic priming without an additional choice element as this allowed us to tap more directly into the processes involved in sentence planning.

The magnitude of the onset latency syntactic priming effects observed in the older adults will be informative about age-related changes in syntactic planning and facilitation that occur during real-time sentence production. While no studies to date have examined age-related effects on onset latency priming, a few studies have investigated age effects on choice syntactic priming. However, this has produced mixed results with two studies finding preserved priming of passives in older adults (Hardy, Messenger, & Maylor, 2017; Hardy, Wheeldon, & Segaert, 2019), while others have not (Heyselaar, Segaert, Walvoort, Kessels, & Hagoort, 2017, footnote 2; Sung, 2015, 2016).^2^ It is therefore difficult to make direct hypotheses about age-related effects on onset latency syntactic priming based on previous evidence. Nevertheless, hypotheses can be made by considering the two-stage competition model in combination with more general models of ageing. The model of Segaert et al. (2016) includes a spreading activation architecture whereby recently processed syntactic structures are activated to an above-baseline level, which contributes to decreased selection and planning speed. However, according to Salthouse’s (1996) general slowing model of ageing, declines in overall processing speed with age can substantially decrease the speed of spreading activation throughout a cognitive or neural network. Similarly, the transmission deficit model postulates that ageing weakens the strength of activation of different units and the connections amongst units, both critical for successful spreading activation (MacKay & Burke, 1990). Applied to syntactic priming, this may mean that when older adults process a prime sentence, the syntactic information relating to the prime does not become available to a central processor quickly or strongly enough to sufficiently excite the representation of the syntactic structure to a level which may influence the speed of syntax selection and planning. If this is the case, we might expect that the magnitude of the onset latency priming effect (i.e., the speed benefit when the syntactic structure is repeated) to be greater for young adults (who possess a faster spreading activation network) compared to older adults (who generally display much slower processing speed; Salthouse, 2004).

## Experiment 1: Method

### Participants

We recruited 50 young adults (36 female) aged 18-25 (*M* = 19.8, *SD* = 1.1) from the University of Birmingham student population and 56 older adults (37 female) aged 64-80 (*M* = 71.8, *SD* = 4.5) from the Patient and Lifespan Cognition Database. Sample sizes were larger than previous studies investigating latency effects of syntactic priming and planning scope (typically 24-34 participants; e.g., Martin, Yan, & Schnur, 2014; Smith & Wheeldon, 2001) and the one previous study that has examined age-related effects in on-line sentence production (15-17 participants per age group; Spieler & Griffin, 2006). All older adults scored above 26 out of 30 (*M* = 27.4; *SD* = 1.3) on the Montreal Cognitive Assessment (Nasreddine et al., 2005), indicating that they were currently experiencing healthy ageing (scores < 26 indicate risk of mild cognitive impairment or dementia; Smith, Gildeh, & Holmes, 2007). All participants were native English speakers with normal or corrected-to-normal vision, and did not report any language disorders. There was no significant difference in education between age groups.^3^ The study was approved by the University of Birmingham Ethical Review Committee and participants provided written informed consent. All participants completed Experiment 1 at an initial test session, followed by Experiment 2 3-7 days later.

### Design

We used a 2 X 2 mixed design with one between-participant variable of age (young vs. older) and one within-participant variable of prime type (syntactically related vs. syntactically unrelated). Hence, there were two experimental task conditions (Figure 1A).

**Figure 1.**
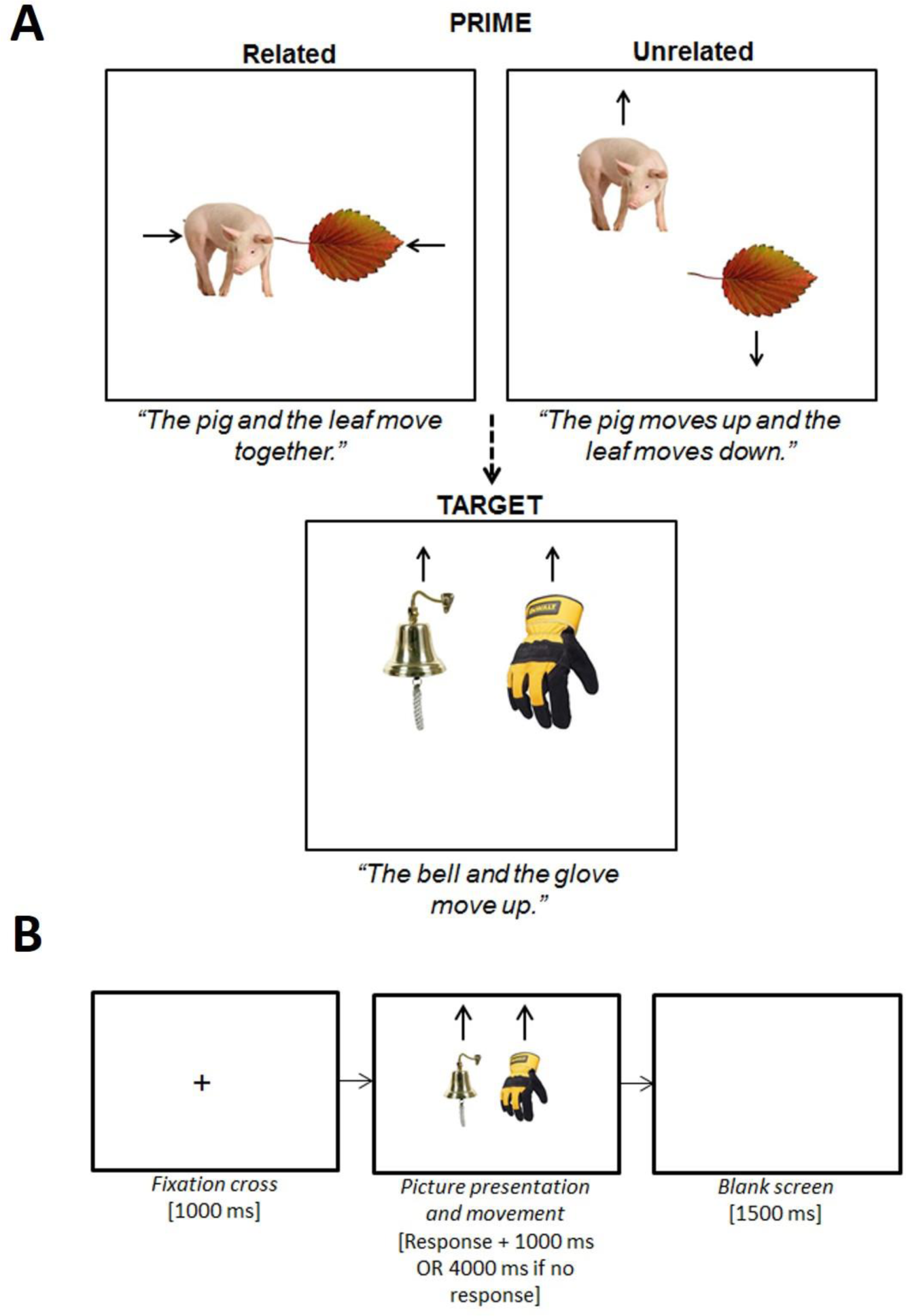
Experiment 1 syntactic priming task design (A) and stimuli presentation events per trial (B). The participant was instructed to begin describing the picture movement as soon as possible using specific sentence types. The stimuli presentation sequence was the same for prime and target trials, and primes were always immediately followed by the corresponding target (i.e., we used a 0-lag delay). Speech latencies on the target trials were recorded from the onset of the pictures to the participant beginning to speak.

### Materials

To create the experimental items, we used 80 simple photographic pictures of everyday concrete objects. All picture names were mono- or disyllabic, and when choosing the stimuli, we took care to ensure that the objects could be identified and named quickly and easily. Close attention to participants’ performance during the practice sessions also indicated that participants did not have issues with picture naming for our specific stimuli. Forty of the pictures were used to create the 40 picture pairs for the target trials; each picture appeared in two different pairs (once each in the left and right position). Using the same constraints, we constructed 40 picture pairs from another 40 pictures for the prime trials. We then paired each target pair with a prime pair to generate 40 experimental items. We ensured that there was no phonological or conceptual overlap between any of the four pictures within each experimental item (this ensured that any effects we observed were related to syntactic processing, and not to semantic or pragmatic features).

The movement of each picture pair was controlled using E-Prime (Schneider, Eschman, & Zuccolotto, 2002). In all target trials, both pictures moved in the same vertical direction (either up or down). Participants were instructed to describe the picture movements from left to right using specific sentences that they were trained on prior to beginning the task; hence, the target trials elicited a coordinate noun phrase (*“the A and the B move up/down”*). In the related prime condition, the pictures moved in opposing horizontal directions which elicited a sentence that was syntactically related to the target trials (*“the C and the D move together*/*apart”*). In the unrelated prime condition, the pictures moved in opposing vertical directions which elicited a sentence that was syntactically unrelated to the target trials (*“the C moves up/down and the D moves down/up”*). We then created two item lists that each contained the same 40 target sentences, but the prime condition matched to each target was rotated such that there were 20 related and 20 unrelated primes per list. Each participant was randomly assigned to one of the two lists and completed 20 experimental items (prime plus target pairs) from each condition (Table 1A). A total of 20 items per experimental condition follows the recommendation of Simmons, Nelson, and Simonsohn (2011) for conducting a well-powered and reliable study.

**Table 1.**
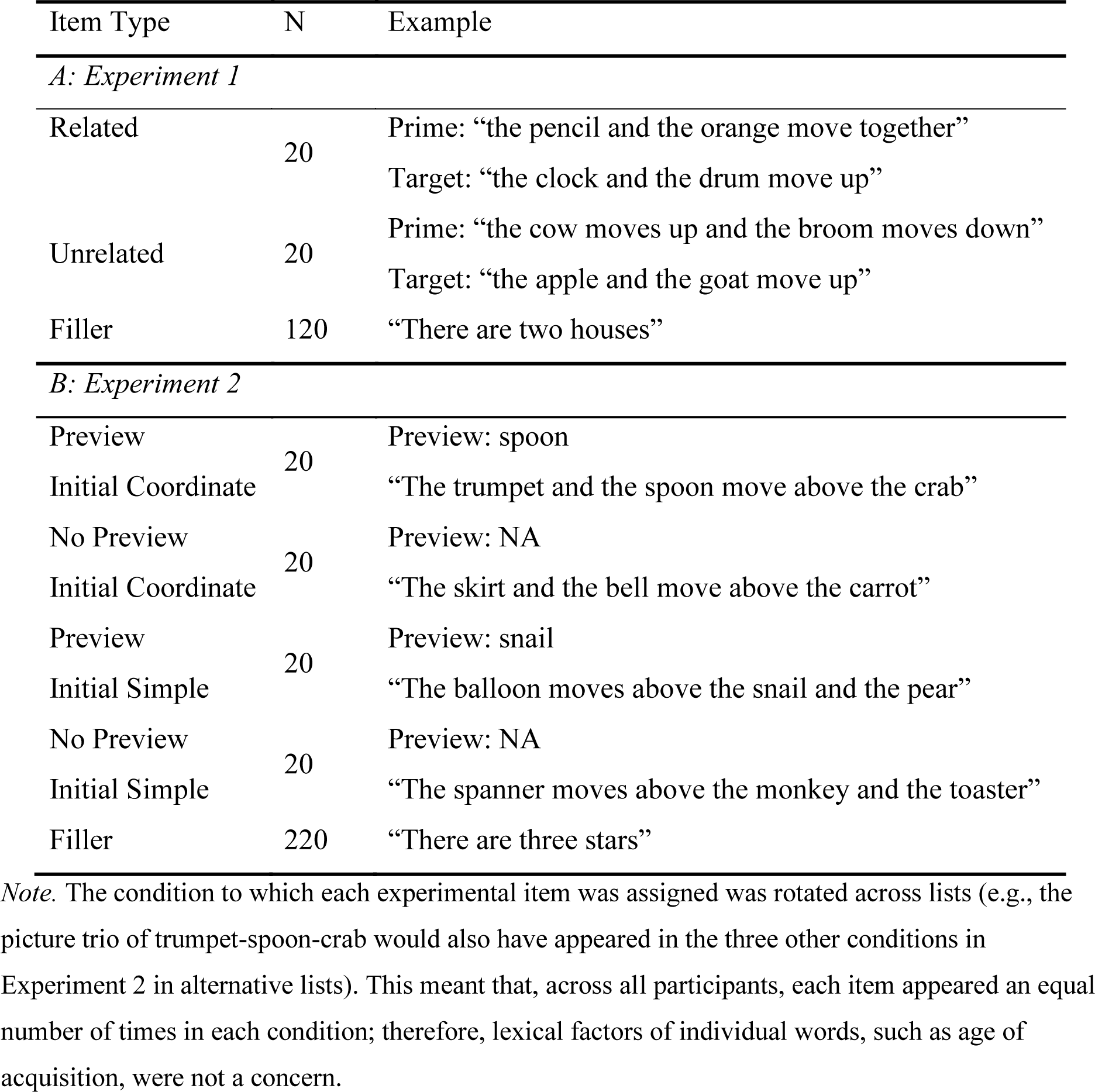
Overview of the different items used in the Experiments 1 and 2. Number of items completed by each participant and example stimuli are provided.

Lastly, we used a further 54 pictures to construct 120 filler trials designed to increase the variety of syntactic structures produced by the participant and to minimise the risk that they would notice the priming manipulation. We created 96 filler trials that elicited phrases such as *“there is an X and a Y”* (no picture movement), *“the Xs move up”* (two repeat pictures move simultaneously), and *“there are no pictures”* (screen is blank). We also created 24 filler trials that elicited phrases that were syntactically similar to the experimental trials; without such ‘decoy’ fillers, experimental trials would always occur in pairs (i.e., prime and corresponding target) which may enable the participant to predict the upcoming movement of a target trial. All 120 fillers were added to each of the two item lists. We then divided each list into four blocks that each contained 5 related experimental items, 5 unrelated experimental items and 30 filler items. The distribution of items within each block was pseudorandomized with the constraint that two experimental items never occurred consecutively. The ordering of the blocks was rotated across participants.

### Procedure

Each participant was tested individually in a sound-attenuating booth facing the screen of a 17 inch *Dell* monitor, in front of which was a *Sony* microphone connected to an amplitude voice key that recorded his/her responses and onset latencies. Figure 1B illustrates the sequence of stimuli presentation per trial. To begin, there were 50 practice trials; the sentences elicited resembled those in the experimental and filler trials and featured all 80 experimental pictures once. If, during the practices, the participant made a lexical error (i.e., used the incorrect picture name) or syntactic error (i.e., used the wrong sentence type), they were corrected by the experimenter. The task then continued until all four experimental blocks had been completed. The experimenter listened from outside the booth via headphones and noted down any errors made by the participant. Errors included: incorrect picture naming (e.g., ‘fish’ instead of ‘shark’); the use of a different sentence structure (e.g., *“the pig moves towards the leaf”* instead of *“the pig and leaf move together”*); and disfluencies, such as pausing and non-lexical errors (e.g., ‘um’).

### Data Preparation and Analyses

We excluded the data of participants whose error rates were above 50% on the experimental trials; this resulted in exclusion of five older adults. Of the 4040 target responses, we excluded trials in which the participant made an error on the corresponding prime (170 (8.5%) of young and 301 (14.7%) of older adult trials). Following Ratcliff’s (1993) recommendation for dealing with reaction time outliers, we also removed trials for which the target onset latency was below 300ms, above 3000ms or more than 2.5SD above/below the participants’ mean per experimental condition (discarding 53 (2.9%) young and 49 (2.8%) older adult trials). All remaining trials were used in the error analyses, but only correct responses (87.4% of trials) were used in onset latency analyses.

All data were analysed in R (R Core Team, 2015) using generalised linear mixed-effects models (*lme4* package; Bates, Mächler, Bolker, & Walker, 2014); this was the most suitable way to analyse the datasets as there were repeated observations for participants and items (Barr, Levy, Scheepers, & Tily, 2013; Jaeger, 2008). We fitted a binomial distribution to the error data as the dependent variable was categorical (correct = 0; incorrect = 1).

Following Lo and Andrews’ (2015) recommendation for analysing continuous speed data, we fitted an inverse Gaussian distribution to the onset latency data with an ‘identity link’ function. This model fit is particularly advantageous when comparing groups with large overall speed differences (i.e., young vs. older) as it eliminates the need for data transformation (i.e., logarithmic or z-scores) while still satisfying the normality assumptions of the generalised linear mixed-effect model (Balota, Aschenbrenner, & Yap, 2013; Lo & Andrews, 2015). For both models, we entered age group (young vs. older) and prime type (related vs. unrelated) as fixed effects. We included random intercepts for participants and items, as well as by-participant and by-item random slopes appropriate for the design. Prior to analysis, the fixed effects were sum-coded and transformed to have a mean of 0 and a range of 1. When a model did not converge with the maximal random effects structure, we simplified the random slopes, removing interactions before main effects in the order of least variance explained until the model converged (Barr et al., 2013).

Given that the effect of ageing was critical to our research question, in the case of non-significant interactions involving age group, we sought to quantify the likelihood of this null effect with additional Bayesian analysis. Using the *BayesFactor* package (Morey & Rouder, 2018), we constructed a full model that included the interaction of interest (H1) and a null model that excluded the interaction (H0); we then calculated the Bayes Factor (BF) as H1/H0. We interpreted the BF values in line with Lee and Wagenmakers’ (2013) classification scheme (see also Schönbrodt & Wagenmakers, 2018). BF values < 0.1 provide ‘strong’ evidence in support of the null (H0) hypothesis, whereas values between 0.1 and 1 are generally deemed inconclusive.

#### Supplementary Measurements

All participants also completed a battery of eight additional measures designed to provide an indicator of their current ability across a variety of cognitive and physical domains. Extensive details about these measurements are available online in the ‘Supplementary Measurements’ section of the OSF repository (https://osf.io/wp7dr/).

## Experiment 1: Results

Figure 2 summarises the target onset latencies and error rates across the two prime conditions for young and older adults.

**Figure 2.**
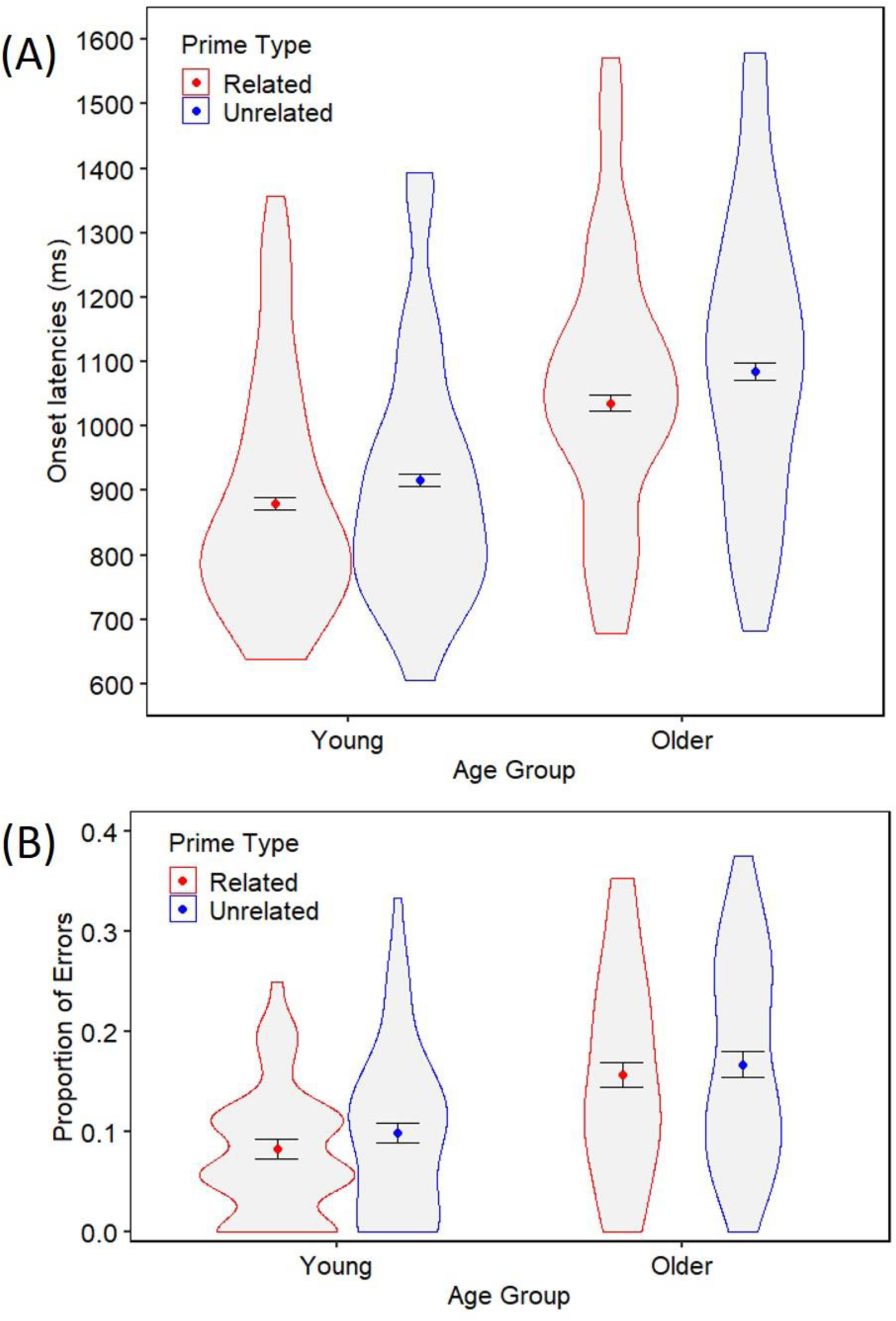
Experiment 1 target onset latencies (A) and errors rates (B) for young and older adults following syntactically related and unrelated primes. The coloured points represent the mean per condition. Error bars denote ±1 the standard error of the mean. Violin spreads represent the distribution of the data across participants.

### Onset Latencies

The best-fitting model of the onset latency data is reported in Table 2A. As expected, older adults were significantly slower than young adults (1060ms vs. 898ms, *p* < .001). There was also a main effect of prime type (*p* < .001), such that target responses were produced significantly quicker following related primes (953ms) than following unrelated primes (994ms), indicating an overall syntactic priming effect of 41ms. Most interestingly, there was no interaction between age group and prime type (*p* = .746), indicating that the onset latency priming effect was similar for young (36ms, 3.9% benefit) and older (49ms, 4.5% benefit) adults. ^4^ Moreover, the additional Bayesian analysis provided ‘strong’ support for the null hypothesis (BF = 0.072) and separate group analyses confirmed that the priming effect was significant for both age groups (Table 2B and 2C).

**Table 2.** Summary of the best-fitted models for the Experiment 1 onset latency data.

| Predictor | Coefficient | SE | *t*-value | *p* |
| --- | --- | --- | --- | --- |
| *A: all data* |  |  |  |  |
| Intercept | 1091.39 | 22.75 | 47.97 | < .001 |
| Prime type | 46.87 | 12.01 | 3.90 | < .001 |
| Age group | -131.40 | 29.24 | -4.49 | < .001 |
| Prime type * Age group | -6.31 | 19.48 | -0.32 | .746 |
| *B: young adults* |  |  |  |  |
| Intercept | 981.89 | 33.63 | 29.19 | < .001 |
| Prime type | 34.59 | 14.22 | 2.43 | .015 |
| *C: older adults* |  |  |  |  |
| Intercept | 1183.26 | 32.93 | 35.94 | < .001 |
| Prime type | 49.11 | 17.10 | 2.87 | .004 |

*Note.* All three models converged with random intercepts for participants and items with additional by-participant and by-item random slopes for the main effects of prime type.

### Error Rates

The best-fitting model of the error data is reported in Table 3A. Although older adults were significantly more error-prone than young adults (16.1% vs. 9.1%, *p* < .001), there was no main effect of prime type (*p* = .369) and no interaction between age group and prime type (*p* = .868; supported by a ‘strong’ BF value 0.060). This suggests that neither young nor older adults’ production of errors on the target trials were affected by the syntactic relatedness of the prime (as was confirmed by separate age groups analyses; Table 3B and 3C).

**Table 3.** Summary of the best-fitted models for the Experiment 1 error data.

| Predictor | Coefficient | SE | Wald Z | *p* |
| --- | --- | --- | --- | --- |
| *A: all data* |  |  |  |  |
| Intercept | -2.34 | 0.16 | -14.69 | < .001 |
| Prime type | -0.14 | 0.15 | -0.90 | .369 |
| Age group | -0.76 | 0.20 | -3.74 | < .001 |
| Prime type * Age group | 0.05 | 0.28 | 0.17 | .868 |
| *B: young adults* |  |  |  |  |
| Intercept | -2.69 | 0.20 | -13.70 | < .001 |
| Prime type | 0.22 | 0.17 | 1.34 | .181 |
| *C: older adults* |  |  |  |  |
| Intercept | -1.96 | 0.18 | -10.90 | < .001 |
| Prime type | 0.10 | 0.16 | 0.61 | .543 |

*Note.* All three models converged with random intercepts for participants and items with additional by-participant and by-item random slopes for the main effects of prime type. The complete dataset model (A) also included a by-item random slope for the main effect of age group.

### Summary

The main findings of Experiment 1 are threefold: (1) older adults were slower and more error-prone when producing sentences compared to young adults; (2) our task produced a reliable latency priming effect on the production of target sentences; and (3) there was no age-related effect in the extent to which the speed of syntax generation benefited from repetition of syntactic structure. Together, this suggests that syntactic facilitation effects on onset latencies are preserved with age.

## Experiment 2: Examining the Effect of Ageing on On-line Planning Scope

In Experiment 1, we demonstrated that syntactic processing in both age groups was facilitated by the repetition of syntactic structure, which in turn benefited the speed of sentence production. This is specifically informative about age-related changes in the processes involved in syntactic facilitation at the planning level of sentence generation, as well as the mechanisms that underlie onset latency syntactic priming. In Experiment 2, we investigated older adults’ sentence generation in unsupported situations in which sentence production is not primed and the speaker must generate a sentence entirely independently. Moreover, we employed a more complex sentence generation task in which participants produced sentences containing multiple phrases of varying length and complexity (this is in contrast to Experiment 1 where the target sentences all consisted of a single coordinate noun phrase). Therefore, within Experiment 2, we aimed to investigate age-related changes in incrementality in sentence production – the scope of sentence planning that occurs prior to articulation onset (Kempen & Hoenkamp, 1987; Levelt, 1989).

A number of studies have demonstrated that speakers do not plan all of what they wish to say before beginning speaking, but instead plan and produce a sentence incrementally in smaller word or phrasal units (see Wheeldon, 2013, for a review). An incremental system is beneficial as it allows for the rapid release of parts of the sentence as soon as planning is complete, reducing the demand for storage in working memory. Previous studies have shown that only a small amount of planning is required prior to speech onset, typically the first phrase (Martin, Crowther, Knight, Tamborello, & Yang, 2010; Martin et al., 2014; Smith & Wheeldon, 1999) or even as little as the first word (Griffin, 2001; Zhao & Yang, 2016).

Moreover, incremental sentence production enables the processing load to be spread across multiple components and time, thereby further reducing demands on cognitive resources (Levelt, 1989; Wheeldon, 2013). One way to investigate the amount of planning in which a speaker engages prior to articulation is with the *planning scope* paradigm, in which picture displays are used to elicit sentences of different syntactic structures and speech onset latencies are used as an on-line measure of advanced planning. For example, Smith and Wheeldon (1999) found that participants took longer to initiate sentences with larger initial coordinate phrases (2a) compared to smaller initial simple phrases (2b). This suggests that planning scope occurs in phrasal units: when the first phrase (defined as the initial conceptual unit that forms a constituent part of a larger syntactic structure) is larger, speakers need longer to plan the syntax and retrieve the second lexical item before speech onset (see also Levelt & Maassen, 1981; Martin, Miller, & Vu, 2004; Wheeldon et al., 2013).

**(2a)** “[the dog and the hat move] above the fork”

**(2b)** “[the dog moves] above the hat and the fork”

Martin et al. (2010, 2014) ruled out an alternative explanation for this effect relating to the visual array (i.e., the grouping of objects moving together) as they found the same phrasal planning scope using stationary picture arrays (e.g., *“the drum and the package are below the squirrel”*). Moreover, the phrasal planning effect cannot be attributed to the fact that, in English, the second content word in the simple initial phrase (always the verb ‘moves’; 2b) may be easier to retrieve than in the coordinate initial phrase (the second lexical item; 2a) as the effect has been demonstrated when the verb changes from trial to trial (Martin et al., 2010), as well as in Japanese, a head-final language in which the subject and the complement take the first two positions in the sentence regardless of initial phrase type (Allum & Wheeldon, 2007, 2009). A phrasal scope of planning has also been demonstrated for other initial phrase structures, such as adjective-noun phrases (e.g., *“the blue frog is next to the blue mug”*; Wagner, Jescheniak, & Schriefers, 2010). Likewise, speakers have been found to take longer to initiate sentences with more complex initial structures (e.g., *“[the river / the large and raging river / the river near their city] empties into the bay…”;* Ferreira, 1991). Nevertheless, the size of speakers’ planning scope is not rigidly fixed and can vary due to multiple factors including ease of syntactic processing (Konopka, 2012; Konopka & Meyer, 2014), task complexity (Ferreira & Swets, 2002; Wagner et al., 2010) and cognitive abilities, such as working memory and production speed (Martin et al., 2004; Slevc, 2011; Swets, Jacovina, & Gerrig, 2014; Wagner et al., 2010). Our interest was in whether the scope of advanced sentence planning is also influenced by healthy ageing.

The ageing process is typically associated with an increase in speech dysfluencies during sentence production, such as the use of non-lexical fillers (‘uh’ or ‘um’), word repetitions and unnatural pauses (Bortfeld, Leon, Bloom, Schober, & Brennan, 2001; Horton, Spieler, & Shriberg, 2010; Kemper, Rash, Kynette, & Norman, 1990). One significant factor that has been proposed to account for this age-related increase in speech dysfluencies is a reduction in the capacity and efficiency of working memory (Abrams & Farrell, 2011; Kemper & Sumner, 2001). This is because verbal working memory is essential for being able to successfully prepare more than one word before beginning articulation (Martin et al., 2004; Slevc, 2011) and for temporarily storing information that is needed for later syntactic processing, such as when producing an embedded clause sentence (Kemper, Kynette, Rash, O’Brien, & Sprott, 1989; Rabaglia & Salthouse, 2011). This suggests that incremental sentence planning processes may become less efficient with age (as the result of declining working memory) or that older adults may adopt different processing strategies when planning a sentence in order to compensate for age-related deficits in working memory. We therefore used the planning scope paradigm to investigate age-related changes in the amount of advanced planning that older speakers engage with prior to articulation.

Based on previous literature, we consider that there are two alternative hypotheses for age-related changes in planning scope. Firstly, a decline in working memory with age may disrupt older adults’ ability to plan sentences with larger initial phrases. Martin et al. (2004) found that an aphasia patient with a semantic working memory deficit displayed a greater phrasal complexity effect than control participants (i.e., a markedly greater difference in the speed of production of larger, compared to smaller, initial phrases), which they attributed to the patient attempting to plan both nouns in the initial phrase, but having difficulty doing so because of deficits at the lexical-semantic level (see also Lee & Thompson, 2011).^5^ Although not as profound as aphasia patients, older adults also experience deficits in working memory (particularly at the verbal level; Bopp & Verhaeghen, 2005). Thus, one hypothesis is that older adults will display a larger phrasal complexity effect than young adults in the planning scope task. Alternatively, to compensate for decline in working memory, older adults may adopt a more extreme word-by-word incremental strategy (i.e., only plan the first word before speech onset regardless of the complexity of the initial phrase). Ferreira and Swets (2002) found that, when time pressure was applied, speakers engaged in significantly less advanced planning, suggesting that incremental planning can be strategically controlled by the speaker. This, combined with the evidence that older adults implement various strategies in other areas of language processing (Altmann & Kemper, 2006; Stine-Morrow et al., 2008), may mean that there is a strategic age-related decrease in the amount of advanced planning that occurs prior to articulation.

In Experiment 2 we further aimed to directly investigate age-related changes in the retrieval of lexical items and their integration into syntactic structures. Lexical retrieval and syntax generation do not rely on the exact same mechanisms, and may even be entirely dissociated (Chang et al., 2000, 2006). Thus, evidence of age effects in syntactic processing does not necessarily mean that age effects will also be observed in lexical processing (or vice-versa). One way to examine lexical processing during sentence production is to incorporate a picture preview element into the planning scope paradigm. Wheeldon et al. (2013) required participants to produce sentences similar to (2a) and (2b), but on some trials there was a preview of one of the upcoming pictures. They found that previewing the second to-be-produced lexical item (*hat* for the examples shown in 2) decreased onset latencies more when it fell within, rather than outside of, the initial phrase. Moreover, Allum and Wheeldon (2009) observed similar latency preview benefits when the preview was presented either in pictorial or written word form, indicating that preview of pictured objects results in lexical access of the name associated with the picture. Together, these findings suggest that the retrieval of lexical items within the first phrase is prioritised prior to speech onset.

Nevertheless, the preview benefit is not reliably maintained when the phrase consisted of three nouns and participants previewed the third lexical item (*“[the drum, the star and the **hat** move] above the crab”*; Wheeldon et al., 2013). Thus, it appears that advanced lexical planning only encompasses a subset of the required nouns and that this does not always align with the scope of syntactic planning.

In Experiment 2 we therefore included a picture preview element within the planning scope task; the magnitude of the preview benefit displayed by older adults will be informative about age-related changes in lexical processing during sentence planning and production.

Young adults’ preferred scope of lexical encoding appeared to be two items (Wheeldon et al., 2013); however, we speculate that older adults’ preferred limit may be less because they have a reduced memory buffer for holding linguistic information (Bopp & Verhaeghen, 2005; Waters & Caplan, 2003). Attempting to retrieve and hold an unmanageable number of lexical items prior to articulation can lead to problems with buffering and maintaining a linearized output (Slevc, 2011; Wheeldon et al., 2013). To overcome this and reduce demands on working memory, older adults may therefore only encode the first lexical item within a phrase prior to articulation; if this is the case, we may expect that, unlike young adults, older adults will not display the preview benefit of the second lexical item even when it falls within the initial phrase.

## Experiment 2: Method

### Participants

The same participants were used as described in Experiment 1.

### Design

We used a 2 X 2 X 2 mixed design with one between-participant variable of age (young vs. older) and two within-participant variables of preview (no preview vs. preview) and initial phrase type (coordinate vs. simple). Hence, there were four experimental task conditions (Figure 3A). Critically, the previewed picture (always of the second upcoming lexical item) fell within the initial phrase in the coordinate condition, but outside of the initial phrase in the simple condition.

**Figure 3.**
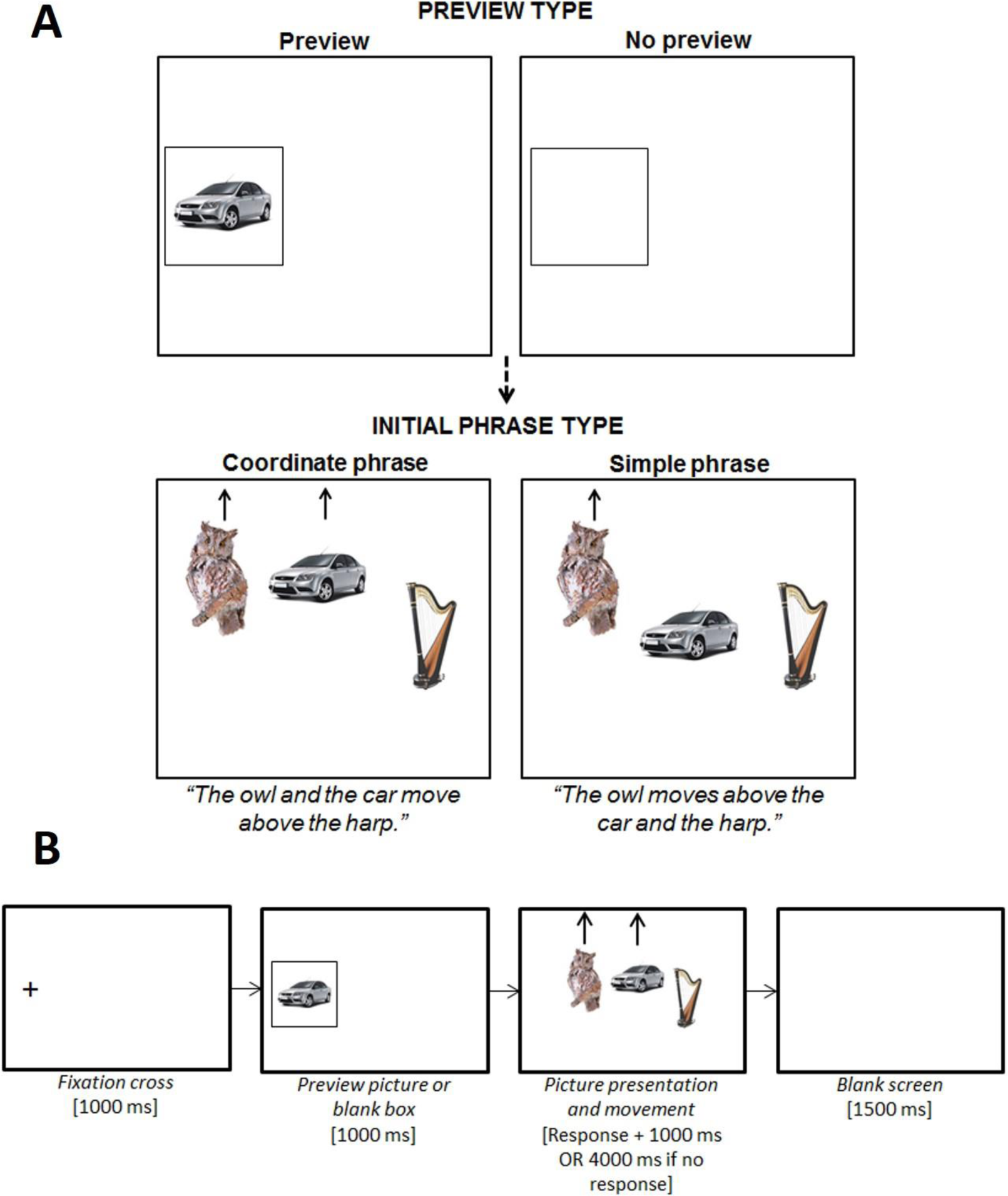
Experiment 2 planning scope design (A) and stimuli presentation events per trial (B). The participant was instructed to pay attention to the preview because it would appear in the upcoming trial, but not to name it aloud. The three pictures then appeared aligned centrally in the horizontal plane (importantly, the leftmost picture did not appear where the preview picture had just been, but in a more right-adjusted position). The participant was instructed to begin describing the picture movement as soon as possible using specific sentence types. Speech latencies were recorded from the onset of the pictures to the participant beginning to speak.

### Materials

To create the experimental items, we used 80 photographic pictures of everyday concrete objects (these were different to those used in Experiment 1, but meet the same criteria). We created 80 experimental items that each consisted of three different pictures that were conceptually and phonologically distinct: each of the 80 pictures appeared in three different experimental items (once in the left, central and right position). As in Experiment 1, the sentence descriptions of the items were elicited by controlling the movement of the pictures (using E-prime) and participants were instructed to describe the picture movements from left to right using specific sentences. In the simple initial phrase conditions, only the left picture moved (either up or down) and the other two pictures remained stationary (*“the A moves above/below the B and the C”*). In the coordinate conditions, both the left and the central picture moved simultaneously (either up or down) and only the right picture remained stationary (*“the A and the B move above/below the C”*). In the preview trials, the preview was always of the central upcoming picture (i.e., object *B*). We created four item lists by evenly rotating the experimental condition assigned to each of the 80 experimental items. Each participant was randomly assigned to one of the four lists and completed 20 experimental items per condition (in line with Simmons et al.’s, 2011, recommendations; Table 1B).

Lastly, we used a further 106 pictures to create 220 filler items designed to prevent the participant from anticipating the location of the preview picture and building expectations to guide their response. The fillers elicited some experimental-type sentences and other sentences that differed from the experimental items in terms of the number of pictures and the type of movement, such as: *“there is an X, a Y and a Z”* (no picture movement); *“the Xs move up”* (three repeat pictures move simultaneously); and “*there are no pictures”*.

Importantly, we also varied the position of the preview pictures within the fillers, such that across all the experimental and filler items each screen position was previewed an equal number of times. All 220 filler items were added to each of the four item lists. We then divided each list into five blocks that each contained 44 fillers and 16 experimental items (4 per condition), and pseudorandomized the order of items using the same constraints as Experiment 1. The ordering of the blocks was rotated across participants.

### Procedure

Each participant was tested using the same equipment set-up described in Experiment 1. Figure 3B illustrates the sequence of stimuli presentation per trial. In the preview trials, the previewed picture was presented for 1000ms: the participant was instructed to pay attention to the preview because it would appear in the upcoming trial, but not to name it aloud. To begin, there were 40 practice trials; the sentences elicited resembled those in the experimental and filler trials and featured all 80 experimental pictures once.

During the practices, the experimenter corrected the participant if they made a lexical or syntactic error. The task then continued until all experimental five blocks had been completed. Using the same criteria described in Experiment 1, the experimenter noted down any errors made by the participant.

### Data Preparation and Analyses

One older adult was excluded from Experiment 2 because of error rates above 50% on the experimental trials. For the 8400 experimental trials, we applied the same onset latency exclusion criteria described in Experiment 1, resulting in the discarding of 124 (3.1%) young and 166 (3.8%) older adult trials. All remaining trials were used in the error analyses, but only correct responses (81.7% of trials) were used in the onset latency analyses.

The data from Experiment 2 were analysed using the same generalised linear mixed-effects modelling methods described in Experiment 1 (a binomial distribution fitted to the error data and an inverse Gaussian distribution fitted to the onset latency data). We entered age group (young vs. older), initial phrase type (coordinate vs. simple) and preview type (no preview vs. preview) into the models as fixed effects. We included random intercepts for participants and items, as well as by-participant and by-item random slopes appropriate for the design. In the case of non-significant interaction involving age group, we used Bayesian analysis to quantify the likelihood of the null effect.

## Experiment 2: Results

Figure 4 summarises the onset latencies and error rates across the four experimental conditions for young and older adults.

**Figure 4.**
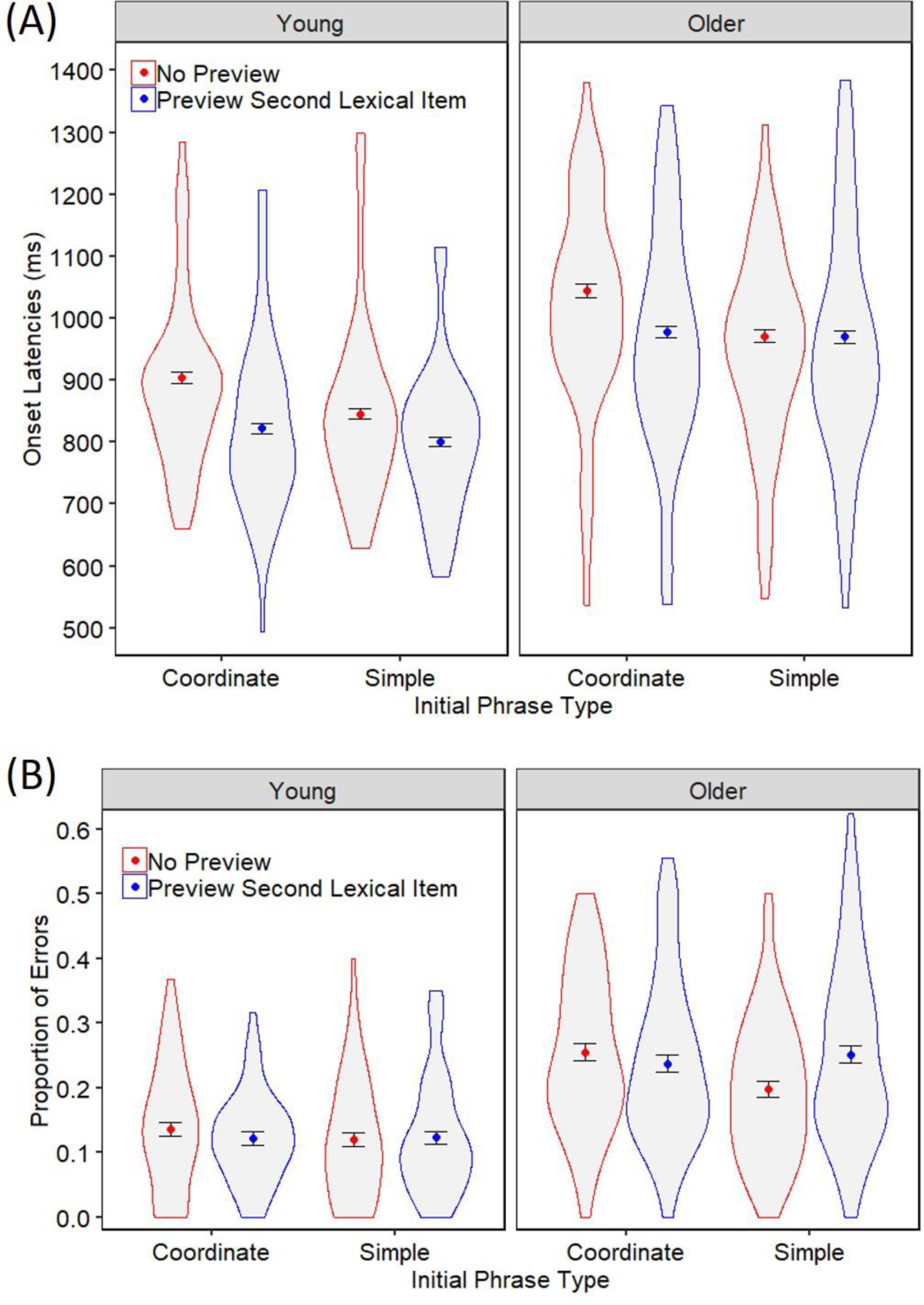
Experiment 2 onset latencies (A) and errors rates (B) for young and older adults when producing sentences within initial coordinate and simple phrases following no preview or a preview of the second upcoming lexical item. The coloured points represent the mean per condition. Error bars denote ±1 the standard error of the mean. Violin spreads represent the distribution of the data across participants.

### Onset Latencies

The best-fitting model of the onset latency data is reported in Table 4A. As in Experiment 1, older adults were significantly slower than young adults (991ms vs. 843ms, *p* < .001). There was a main effect of initial phrase type, such that sentences with initial simple phrases were produced significantly quicker than sentences with initial coordinate phrases (895ms vs. 935ms, *p* < .001), indicating an overall phrasal planning effect of 40ms (4.5%). Furthermore, the interaction between initial phrase type and age group was not significant (*p* =. 994), indicating that the incremental planning effect was unaffected by healthy ageing, as was supported by an ‘extremely strong’ Bayes Factor (BF) value (0.004). Indeed, separate age group analyses confirmed that the phrasal planning effect was highly significant for both young (40ms, 4.6% benefit, *p* < .001) and older (41ms, 4.0% benefit, *p* < .001) adults.

**Table 4.** Summary of the best-fitted models for the Experiment 2 onset latency data.

| Predictor | Coefficient | SE | Wald Z | *p* |
| --- | --- | --- | --- | --- |
| *A: All data* |  |  |  |  |
| Intercept | 1008.53 | 14.44 | 69.86 | < .001 |
| Preview | -57.60 | 8.73 | -6.60 | < .001 |
| Initial phrase type | -43.33 | 6.91 | -6.27 | < .001 |
| Age group | -132.07 | 16.07 | -8.22 | < .001 |
| Preview * Initial phrase type | 51.01 | 8.77 | 5.82 | < .001 |
| Preview * Age group | -25.26 | 12.03 | -2.10 | .036 |
| Initial phrase type * Age group | 0.29 | 11.52 | 0.03 | .980 |
| Preview * Initial phrase type * Age group | -22.09 | 14.25 | -1.55 | .121 |
| *B: Young Adults* |  |  |  |  |
| Intercept | 911.75 | 28.52 | 31.97 | < .001 |
| Preview | -67.95 | 16.36 | -4.15 | < .001 |
| Initial phrase type | -44.74 | 9.49 | -4.72 | < .001 |
| Preview * Initial phrase type | 40.68 | 16.98 | 2.40 | .017 |
| *C: Older Adults* |  |  |  |  |
| Intercept | 1109.87 | 21.08 | 52.66 | < .001 |
| Preview | -43.59 | 18.05 | -2.41 | .016 |
| Initial phrase type | -39.42 | 11.66 | -3.38 | < .001 |
| Preview * Initial phrase type | 60.56 | 14.26 | 4.25 | < .001 |

*Note.* All three models converged with random intercepts for participants and items with additional by-participant random slopes for the main effects of preview and initial phrase type, and a by-item random slope for the main effect of preview.

The analyses further revealed a main effect of preview, such that sentences were produced significantly quicker following preview of the second upcoming lexical item compared to no preview (890ms vs. 939ms, *p* < .001), indicating an overall preview benefit of 49ms (5.5%). Interestingly, there was a significant interaction between preview and age group (*p* = .036), such that the preview benefit was larger for young (62ms, 7.6%), compared to older (33ms, 3.4%), adults. Moreover, there was a significant interaction between preview and initial phrase type (*p* < .001): the overall preview benefit was significantly greater when the preview picture fell within the initial phrase (coordinate condition; 74ms, 7.6%) compared when it fell outside of it (simple condition; 26ms, 2.9%).

Although the three-way interaction between preview, initial phrase type and age group did not reach significance (*p* = .121), the Bayesian analysis provided inconclusive evidence in support of the null hypothesis (BF = 0.141). Moreover, separate age group analyses (Tables 4B and 4C) suggest that the significant difference in the preview effect for young and older adults may have been driven by more complex effects at the phrase level. Both young (*p* = .017) and older (*p* < .001) adults showed a significant interaction between phrase type and preview; however, this may represent different pattern of effects for each age group (see Figure 4A). Further post-hoc pairwise comparisons revealed that for young adults, there was a significant benefit of preview in both the coordinate (81ms (8.9%), χ^2^(1) = 18.20, *p* < .001) and simple (45ms (5.3%), χ^2^(1) = 9.03, *p* = .002) phrase conditions, although the magnitude of the effect was distinctly larger when the preview fell within the initial phrase.^6^ By contrast, the difference in onset latencies between preview conditions was only significant for the older adults when it fell within the initial phrase (67ms (6.4%) preview benefit; χ^2^(1) = 15.18, *p* < .001), but not outside of it (2ms (0.2%) preview benefit; χ^2^(1) = 0.45, *p* = .502).

### Error Rates

The best-fitting model of the error data is reported in Table 5A. As in Experiment 1, older adults were significantly more error-prone than young adults (23.5% vs. 12.5%, *p* < .001). While there were no main effects of preview (*p* = .308) or initial phrase type (*p* = .097), there was a significant interaction between the two variables (*p* = .040): the presence of the preview resulted in a 1.6% decrease in participants’ errors when producing sentences with initial coordinate phrases, but a 2.9% increase in errors when producing sentences with initial simple phrases. There was no significant interaction between phrase type and age group (*p* = .747; supported by an ‘extremely strong’ BF value of 0.009) or between preview and age group (*p* = .292; supported by an ‘very strong’ BF value of 0.017). There was also no interaction between preview, initial phrase type and age group (*p* = .295); however, in the case of this 3-way interaction, the Bayesian analysis provided inconclusive support for the null hypothesis (BF = 0.190).

**Table 5.** Summary of the best-fitted models for the Experiment 2 error data.

| Predictor | Coefficient | SE | Wald Z | *p* |
| --- | --- | --- | --- | --- |
| *A: All data* |  |  |  |  |
| Intercept | 2.02 | 0.15 | 13.62 | < .001 |
| Preview | -0.07 | 0.07 | -1.02 | .308 |
| Initial phrase type | 0.12 | 0.07 | 1.66 | .097 |
| Age group | 0.89 | 0.16 | 5.70 | < .001 |
| Preview * Initial phrase type | -0.28 | 0.14 | -2.06 | .040 |
| Preview * Age group | 0.14 | 0.14 | 1.05 | .292 |
| Initial phrase type * Age group | -0.04 | 0.14 | -0.32 | .747 |
| Preview * Initial phrase type * Age group | 0.29 | 0.27 | 1.07 | .285 |
| *B: Young Adults* |  |  |  |  |
| Intercept | -2.50 | 0.17 | -14.65 | < .001 |
| Preview | 0.06 | 0.12 | 0.51 | .607 |
| Initial phrase type | -0.14 | 0.12 | -1.16 | .245 |
| Preview * Initial phrase type | 0.20 | 0.21 | 0.93 | .352 |
| *C: Older Adults* |  |  |  |  |
| Intercept | -1.59 | 0.17 | -9.18 | < .001 |
| Preview | 0.12 | 0.09 | 1.42 | .154 |
| Initial phrase type | -0.12 | 0.09 | -1.39 | .163 |
| Preview * Initial phrase type | 0.41 | 0.17 | 2.41 | .016 |

*Note.* All three models converged with random intercepts for participants and items with additional by-participant random slopes for the main effects of preview and initial phrase type. The complete dataset model (A) also included a by-item random slope for the main effect of age group.

Further insight into possible age-related effects may be gleamed from separate age group analyses (Tables 5B and 5C). This revealed that the interaction between preview and initial phrase type remained significant for older adults (*p* = .016), but not for young adults (*p* = .352). This suggests that young adults’ error rates were fairly stable across conditions, whereas the proportion of errors produced by older adults differed between phrase types dependent on whether the preview was present. Further post-hoc comparisons revealed that there was no effect of preview on older adults’ error rates when it fell within the initial phrase (coordinate condition; χ^2^(1) = 0.32, *p* = .570), but that the presence of the preview caused a significant 5.3% increase in errors when it fell outside of the initial phrase (simple condition; χ^2^(1) = 8.35, *p* = .003).

### Summary

The main findings of Experiment 2 can be summarised as follows: (1) as in Experiment 1, older adults were slower and more error-prone than young adults; (2) our task elicited a reliable phrasal planning scope effect that was unaffected by healthy ageing; and (3) while young adults’ displayed speed benefit of preview in both phrase conditions, older adults only benefited when the preview fell within the initial phrase and produced significantly more errors when the previewed lexical item fell within the second phrase.

Together, this suggests that there were age group differences in lexical processing during sentence planning which only emerged when the preview fell outside of the initial phrase. It should be noted, however, that a potential caveat of our findings is that we did not find a higher-order interaction between age group, preview and initial phrase type (and the following Bayesian analysis did not provide conclusive support for either the null or alternative hypothesis). As such, our post-hoc analyses should be considered somewhat exploratory in nature and we emphasise the need for replication in future studies.

Nonetheless, we do still consider our findings to provide a valuable and interesting insight into possible age group differences. Indeed, Fiedler, Kutzner, and Krueger (2012) argue that it is important to rigorously explore all possible findings within a dataset, even when they are accompanied by some apparently null results, in order to prevent against the risk of a false negative (the discovery of a false null result) (see also Wei, Carroll, Harden, & Wu, 2012).

## General Discussion

Using two on-line experiments, we investigated age-related changes in the syntactic and lexical processes involved in sentence generation. In Experiment 1, both young and older adults produced target sentences quicker following syntactically related primes, demonstrating that speed benefits of syntactic priming are preserved with age, despite older adults’ slower and more error-prone production. In Experiment 2, both young and older adults initiated sentences quicker with smaller, compared to larger, initial phrases, suggesting that planning scope, at least at the syntactic level, is unaffected by healthy ageing. Evidence of age-related differences did emerge, however, in the preview conditions, such that young adults displayed a significantly larger preview benefit than older adults (quicker to initiate sentences when there was a preview of an upcoming lexical item). Moreover, post-hoc analyses demonstrated that, while young adults displayed speed benefits of preview when the pictured word fell both within and outside the initial phrase, older adults only displayed speed benefits from the previewed picture when it fell within the initial phrase, and preview outside of the initial phrase caused them to become more error-prone. This suggests that age differences may exist in the flexibility of lexical retrieval during sentence planning and in the ability to integrate lexical information into syntactic structures. Taking both experiments together, our study therefore suggests age-related effects of lexical, but not syntactic, processes on the speed and accuracy of sentence production.

Our robust finding of equal onset latency priming in both age groups in Experiment 1 (supported by both traditional null hypothesis testing and Bayesian analysis) provides the first evidence that syntactic facilitation effects are preserved with age in a task specifically designed to tap into the processes involved in the planning stage of sentence production.

Applied to Segaert’s et al. (2016) two-stage competition model, this suggests that older adults maintain the ability to quickly and efficiently generate previously activated syntactic structures. This is somewhat contrary to our initial hypothesis that overall decline in processing and transmission speed with age would result in decreases in the spreading activation architecture that supports syntactic priming (MacKay & Burke, 1990; Salthouse, 1996). Instead, the slowing associated with ageing might not affect all cognitive networks equally (Fisher, Duffy, & Katsikopoulos, 2000; Fisk, Fisher, & Rogers, 1992). Thus, despite general slowing elsewhere, older adults appear to maintain sufficient cognitive resources to support successful syntactic priming. This is consistent with the evidence of priming benefits in older adults in other areas of language processing, such as semantic priming (Burke, White, & Diaz, 1987; Laver & Burke, 1993) and morphological priming of both regularly-inflected verbs and transparent compounds (Clahsen & Reifegerste, 2017; Duñabeitia, Marín, Avilés, Perea, & Carreiras, 2009; Reifegerste, Elin, & Clahsen, 2018). Together with our findings, this indicates that models of language and ageing should account for the effects of process-specific, rather than general, cognitive slowing (see Laver & Burke, 1993, for a more extensive discussion).

Nevertheless, it is important to consider that we found evidence of preserved latency priming effects in older adults in a task in which the demands were relatively low: participants only needed to dedicate minimal cognitive resources to syntactic selection (because we removed the choice element) and we did not manipulate the ease of lexical encoding. According to Peelle (2019), the relationship between cognitive supply and task demands would still therefore be balanced in favour of good behavioural performance in older adults, despite likely declines in overall cognitive capacity. It therefore remains unclear whether latency priming effects would continue to be observed in older adults in a task in which demands are increased (e.g., by manipulating the codability of the nouns). Moreover, the consideration of task demands vs. cognitive supply may also be necessary for clarifying the mixed findings within the existing choice syntactic priming and ageing literature (Hardy et al., 2017, 2019; Heyselaar et al., 2017; Sung, 2015, 2016). There are minimal methodological differences between the various studies (e.g., all used a picture description production task); however, it remains possible that differences in the characteristics of the samples, such as education level and native language use, may have resulted in differences in processing efficiency between the older adult groups, leading to different behavioural findings between studies (Peelle, 2019). Unfortunately, this information is unavailable for previous studies, meaning such a comparison is not possible. This highlights why it is important for future research to collect individual differences data, as well as age group information, when investigating what determines latency and choice syntactic priming.

Turning now to the findings of Experiment 2, the pattern we observed in the onset latencies is similarly consistent with an age-related preservation of syntactic processing skills as we found robust evidence of a phrasal scope of planning in both age groups: speakers took longer to initiate sentences with larger initial phrases. This replicates previous research in young adults (e.g., Martin et al., 2010, 2014; Smith & Wheeldon, 1999), and suggests that both age groups prioritised the generation of syntax within the first phrase prior to articulation. It is notable that older speakers did not experience disproportionate difficulty in planning the larger initial phrases (as has been observed in aphasia patients; Martin et al., 2004), indicating that, although ageing is associated with decline in general cognitive function, this is not substantial enough to cause age-related deficits in incremental sentence production. Moreover, our findings demonstrate that older adults do not actively engage in a more extreme word-by-word planning strategy (if this was the case, latencies would have been similar for simple and coordinate initial phrases), further suggesting that older adults maintain sufficient cognitive capacity to support the planning of an initial phrase containing at least two nouns. Spieler and Griffin (2006) also found no differences in the sentence planning strategy used by young and older adults; however, they found that both age groups planned in single word, not phrasal, units. This apparent contrast to our findings can likely be explained by the different measurements used; specifically, while our use of onset latency measures provided insight into the preparation time before sentence articulation, Spieler and Griffin’s (2006) use of eye-tracking focused more on the gaze shifts that occur during articulation and which are tightly locked to individual word onset. Nevertheless, both findings indicate that there are minimal age group differences in on-line syntactic processing, as has been found in other studies in which participants are presented with different words on screen and asked to formulate a sentence (Altmann & Kemper, 2006; Davidson et al., 2003).

An important point to make, however, is that the minimal age group differences we observed in syntactic planning do not necessarily mean that young and older adults were engaging the exact same cognitive networks when performing the task. While young adults may be predominantly relying on activity in the left anterior temporal lobe and the left inferior frontal gyrus to support incremental sentence planning (Brennan & Pylkkänen, 2017; Ohta, Fukui, & Sakai, 2013; Snijders et al., 2008; Uddén et al., 2019), older adults may be recruiting additional areas outside of the core language network to support performance (in the same way as has been observed for other aspects of language processing; Peelle, Troiani, Wingfield, & Grossman, 2010; Wingfield & Grossman, 2006). Further work is therefore needed to fully understand the age-related changes in the neural networks that underlie incremental sentence planning. Indeed, evidence of age group differences did emerge due to the picture preview manipulation, suggesting that young and older adults may be adopting different strategies relating to lexical processing, a finding we turn to next.

In Experiment 2, half of the experimental trials were preceded by a picture of the upcoming second lexical item. Overall both age groups were quicker to initiate sentences when there was a preview compared to no preview; however, the magnitude of the preview benefit was significantly greater for young, compared to older, adults. This suggests possible age-related effects in speakers’ abilities to incorporate previewed lexical information into their sentence planning. Moreover, further post-hoc analyses suggest that age group differences in the preview benefit may have been driven by more complex effects at the phrase level. To first consider when the previewed picture fell within the initial phrase (*“[the owl and the **car** move] above the harp”*), we found that both young and older adults were quicker to initiate the sentence when there was a preview, compared to no preview, suggesting that the prior retrieval of the lexical item was significantly benefiting their sentence planning at the lexical encoding level (Allum & Wheeldon, 2009; Wheeldon et al., 2013). However, to now consider when the previewed picture fell outside of the initial phrase (*“[the owl moves] above the **car** and the harp”*), some interesting age group differences did emerge in participants’ onset latencies and error rates. While young adults continued to display speed benefit of preview outside the second phrase (albeit to a lesser extent than when it fell within the initial phrase), older adults did not display a speed preview benefit when it appeared within the second phrase, and the presence of the picture preview outside their preferred phrasal planning scope caused them to become significantly more error-prone. Importantly, this increase in error rates (including non-lexical dysfluencies) for the older adults is unlikely to relate to specific issues with picture naming and syntax selection due to the large number of practices completed prior to the experimental task (during which the experimenter corrected the participant if they used an incorrect picture name or sentence type), but more due to disruption during the sentence planning process. Taken together, the onset latency and error data therefore suggest that, unlike young adults, older adults did not benefit from this early access to lexical information and that, instead, this premature availability had a disruptive effect on their overall fluency.

One explanation for this age group difference relates to age-related differences in the flexibility of the sentence planning process. The fact that young adults displayed significant preview benefits in both phrase conditions, but to a greater extent when the preview fell within the initial phrase, suggests that they prioritised the retrieval of lexical items within the first phrase prior to articulation, but they were also able to successfully manage the early activation of lexical items outside of their usual phrasal planning scope to benefit their overall speed of sentence production. This evidence of adaptability within young adults’ planning scope adds to the growing evidence that planning scope is flexible and can be influenced by the ease of syntactic and lexical processing (Konopka, 2012; Konopka & Meyer, 2014; van de Velde & Meyer, 2014). By contrast, older adults’ planning scope appears to be a lot more rigidly fixed to phrasal boundaries such that they are less adaptable when it comes to integrating new lexical information into syntactic structures. Indeed, older adults show less parafoveal preview effects across syntactic pauses during sentence comprehension than young adults, suggesting an age-related segmentation strategy designed to aid syntactic processing (Payne & Stine-Morrow, 2012, 2014; Stine-Morrow & Payne, 2016). This segmentation strategy may also apply to older speakers’ sentence production; specifically, in an attempt to decrease processing demands, older adults may strategically choose to only attend to lexical information when it is relevant (i.e., only when it is contained within the next to-be-produced phrase). Thus, older adults are less able to successfully incorporate lexical information outside of the initial phrase into their sentence planning. This contrast between the flexible sentence planning approach observed in the young adults and the rigid approach in the older adults further highlights how apparently similar behaviour in both age groups (i.e., both displayed a phrasal scope of planning) may be supported by different cognitive networks and processing strategies.

A second explanation for the age-related difference in lexical processing that we observed involves the executive control required to successfully manage the premature access to lexical information. During the picture preview, participants automatically access some lexical information about the pictured item which would be stored in their working memory. Given that young and older speakers displayed preview benefits within the first phrase, we consider it likely that participants had sufficient time to access the lemma corresponding to the picture name. Critically, participants would have done this for all preview pictures since the syntactic structure and position of the previewed lexical item in the upcoming trial was unpredictable (due to the use of lots of filler items and stringent counter-balancing).

However, if the previewed lexical item does not appear in the first phrase, participants must temporarily inhibit this information in order to prevent it from interfering with the retrieval of the first (unpreviewed) lexical item and planning of the initial phrase. Therefore, when the previewed lexical item falls outside of the initial phrase, there is increased demand on the cognitive resources, in particular inhibitory control, that support the processes involved in maintaining a linearized output. Young adults appear to be very good at coping with this increased demand as they even benefit from the preview information when it is required in the planning of the second phrase.

By contrast, older adults show no speed benefits of the previewed picture when it fell within the second phrase, and instead the presence of the preview caused them to become significantly more error-prone. Theoretical accounts propose that ageing weakens the inhibitory processes that are responsible for regulating what information enters and leaves working memory (Hasher, Lustig, & Zacks, 2007; Hasher & Zacks, 1988). Specifically, if older adults are less able to engage the required level of inhibitory control, the balance between processing efficiency and task demands will move to favour the latter, resulting in increased inference effects and poorer behavioural performance (Peelle, 2019). Indeed, deficits in inhibitory control have been used to explain other age effects on language processing, such as older adults having increased difficulty ignoring distracting or irrelevant information during speech comprehension and production (Britt, Ferrara, & Mirman, 2016; Sommers & Danielson, 1999; Tun, O’Kane, & Wingfield, 2002). Deficits in inhibitory control may therefore offer a possible explanation for our findings as, if the older adults were less able to inhibit irrelevant lexical information during the planning of the first phrase, this would lead to increased problems with formulating a linearized output, resulting in increased errors. This executive control interpretation of the age effects that we observed may be considered in parallel with our previous interpretation relating to the flexibility of sentence planning since efficient verbal working memory and inhibitory control skills are essential for being able to successfully plan and produce multi-word utterances (Engelhardt, Corley, Nigg, & Ferreira, 2010; Martin et al., 2004; Slevc, 2011). Nonetheless, without evidence of a direct link between participants’ task performance and individual measures of inhibitory control, our executive control explanation remains somewhat speculative. Within the Supplementary Measurements of this study (available online on the OSF: https://osf.io/wp7dr/), we did include eight individual difference measures, such as inhibition, as additional predictors in the two experimental sentence production tasks. However, this did not produce any notable results, something which we likely attribute to our use of a single measurement per construct and the inherent difficulties involved in measuring individual differences within a factorial design (see Supplementary Measurements for a more in-depth discussion). Further work, employing a large battery of inhibition measures, is therefore required to test more directly whether there is a relationship between inhibitory control and lexical planning in healthy ageing.

In summary, our study is the first to examine age-related changes in syntactic and lexical processing during sentence production using on-line onset latency measures. Specifically, our study provides evidence for the age-related preservation of syntactic processing (as evident in the syntactic priming and phrasal planning scope effects we observed in both age groups), but increased difficulty with lexical retrieval and integration with age (older adults displayed less benefits of preview, particularly when the previewed lexical item fell outside of the initial phrase). We attribute this apparent age-related decline in lexical processing to a decline in the flexibility of sentence planning processes, in particular in speakers’ ability to incorporate novel lexical information into their sentence planning. This may be related to older speakers’ stronger preference for segmentation at phrasal boundaries when planning a sentence (a strategic approach designed to minimise processing demands) and/or to a decline in executive control, making older speakers less able to cope with premature lexical activation beyond the first phrase. Our findings should be considered in parallel with off-line studies of language and ageing in order to gain a more complete picture of language processing in old age, in terms of which processes are preserved and which decline.

## Author Note

### Acknowledgements

We gratefully acknowledge the help of Denise Clissett, the coordinator of the Patient and Lifespan Cognition Database. This manuscript has been released as a Pre-Print at *BioRxiv* (https://www.biorxiv.org/content/10.1101/327304v5).

### Funding

This research was partly supported by an Economic and Social Research Council (ESRC) studentship awarded to SMH from the University of Birmingham Doctoral Training Centre (grant number: ES/J50001X/1).

### Author Contributions

SMH, KS and LW designed the study. SMH collected and analysed the data, and wrote the manuscript. All authors revised the manuscript.

### Data Availability

The complete datasets and supplementary measurements of the study are provided online on the Open Science Framework (https://osf.io/wp7dr/).

## Footnotes

1 We note other studies have employed on-line measures to investigate age-related differences at the single word level (see Mortensen, Meyer, & Humphreys, 2006, for a review). While producing single words requires the retrieval of lexical information, it does not require the incorporation of the lexical items into a syntactic structure for sentence production (Levelt, 1989). Thus, it is difficult to apply these single word findings to age-related effects on sentence production (Kavé & Goral, 2017).

2 Note, some other studies tested non-young adults as controls for clinical patients; however, the samples are small and the age ranges are large. While Ferreira, Bock, Wilson, and Cohen, (2008, *n* = 4 aged 50-58) and Cho-Reyes, Mack, and Thompson (2016; *n* = 13 aged 33-76) found evidence of choice syntactic priming in controls, Hartsuiker and Kolk (1998; *n* = 12 aged ∼28-67) did not.

3 Education was scored according to the International Standard Classification of Education (United Nations, 2011), which classifies education on a scale of 0 (pre-primary school) to 8 (university doctorate). There was no significant difference in scores between young (*M* = 6.0, *SD* = 0.1) and older (*M* = 5.8, *SD* = 1.3) adults, *t*(104) = 1.36, *p* = .178. A score of 6.0 indicates engagement in formal education to an undergraduate bachelor level (approximately equal to 17 years).

4 Percentage priming benefit = (Unrelated – Related) / Unrelated

5 We note that both of these studies did include non-young adults as controls for aphasia patients; however, the sample are small and the age ranges large (Lee & Thompson, 2011, *n* = 9 aged 48-73; Martin et al., 2004, *n* = 10 aged ∼55-66), making it difficult to draw any firm conclusions about age-related effects on incremental sentence planning.

6 For the post-hoc analyses, we used the *‘testInteractions’* function in the *phia* package (de Rosario-Martinez, 2015a, 2015b) which allows for the direct comparison of contrasts specified within an existing mixed-effect model. Importantly, the *‘testInteractions’* function corrects *p* values for multiple comparisons using the Holm-Bonferroni method (adjusts the criteria of each individual hypothesis), thereby reducing the risk of discovering a false positive result (Aickin & Gensler, 1996; Holm, 1979).

